# N-Terminal Deleted Isoforms of E3 Ligase RNF220 (Isoform 4) Are Ubiquitously Expressed and Required for Mouse Muscle Differentiation

**DOI:** 10.1101/2025.04.07.647521

**Authors:** SeokGyeong Choi, Donald J. Wolfgeher, Jeewon Kim, Young-Hyun Go, Hyuk-Jin Cha, Gyu-Un Bae, Stephen J. Kron, Woo-Young Kim

**Author notes:** Correspondence should be addressed to Stephen J. Kron and Woo-Young Kim.

## Abstract

Four isoform peptides of the novel E3 ligase RNF220 have been identified in humans. However, all of previous studies have predominantly focused on isoform 1, which consists of 566 amino acids (aa). Here, we show that a shorter isoform, isoform 4 (308 aa), lacking most of the N-terminus, is the predominant and ubiquitously expressed variant that warrants functional investigation. Both isoform 1 and isoform 4 are expressed in the brain; however, isoform 4 is the major isoform expressed in all other tissues in mice. Consistently, H3K4me3 ChIP-seq data from ENCODE reveal that the transcription start site for isoform 4 demonstrates broader and stronger activity across human tissues than that of isoform 1. Isoform 4 produces two peptides (4a and 4b) through alternative translation initiation, with isoform 4b displaying distinct subcellular localization and subnuclear structures. Notably, during embryonic stem cell differentiation into neural stem cells, isoform 1 expression increases, whereas isoform 4 expression decreases. In murine myoblasts, isoform 4 is the sole expressed isoform and is required for MyoD and myogenin expression, as well as for muscle differentiation. Our findings highlight isoform 4 as the ubiquitously and highly expressed variant, likely playing a fundamental role across tissues while exhibiting functional differences from isoform 1. These results emphasize the critical importance of isoform 4 in future studies investigating the biological functions of RNF220.

## Introduction

Ring finger protein 220 (RNF220) is an E3 ubiquitin ligase involved in neural development and synaptic transmission (Ma et al. 2019; Song, et al. 2020; Ma, An, et al. 2020; Wang, et al. 2022; Ma, et al. 2022). Furthermore, it has roles in carcinogenesis and immunity (Pan, et al. 2021; Yan, et al. 2021; Guo, et al. 2021; Deng, et al. 2023).

RNF220, located on chromosome 1p34.1 (NCBI Gene ID: 55182), produces four isoforms using five transcription start sites and alternative splicing (**Fig. 1a**). Shorter isoforms 2 and 4 lack N-terminal regions present in longer isoforms 1 and 3. The protein isoforms from the same gene may mediate unique or even opposing biological functions. As seen with *LEF1* (de Klerk and ‘t Hoen. 2015), *MYC* (Kubickova, et al. 2023), *STAT3* (Aigner, et al. 2019), and *RUNX1* (Davuluri, et al. 2008), RNF220 isoforms may also have distinct biological functions. However, most studies on RNF220 have focused on the ‘full-length’ 63 kDa isoform 1, in gain-of-function and loss-of-function experiments. Widely used RNF220 knockout mice, developed by deleting exon 2 (Ma et al. 2019), may retain isoforms 2 and 4. Should these short isoforms remain expressed in this model, they may be obscuring as-yet unappreciated physiological roles for RNF220.

**Figure 1.**
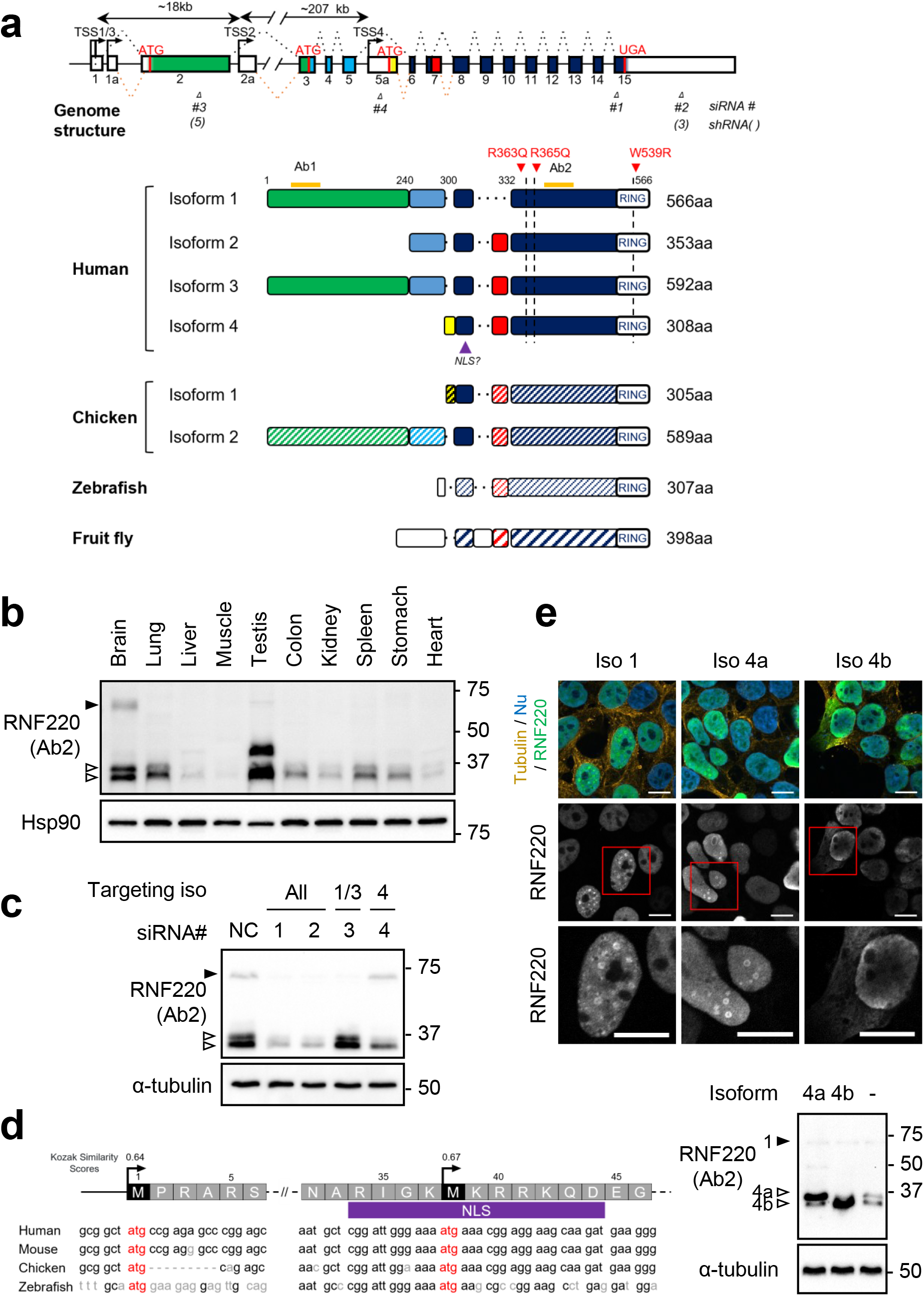
Comparative Analysis of RNF220 Isoforms: Structure, Expression, and Subcellular Localization. (a) Exon structure of human RNF220 isoform 1 mRNA (NM_018150.4) and other RNF220 isoforms and schematic diagram of RNF220 isoforms in human, chicken, zebrafish, and fruit fly based on their amino acid sequence similarity. The length of each exon and protein domain is proportional to its actual length. Global alignment was performed using the Needleman-Wunsch algorithm. (b) Comparison of RNF220 isoform expression in various mouse tissues by western blot (9-week-old, male). (c) Identification of each isoform in HEK293 with siRNAs targeting isoform-specific regions illustrated in (a). (d) Amino acid sequence and cDNA alignment of the RNF220 isoform 4 N-terminal region across species. The annotated AUG start codon and the 2ndary translation start are indicated with their Kozak similarity score (Gleason, et al. 2022). Nuclear localization signal (NLS) predicted by cNLS Mapper (Kosugi, et al. 2009) is marked. Identification of isoform 4a or 4b was performed using HEK293 cells transfected with the corresponding cDNAs. (e) Immunostaining of RNF220 isoform 1, 4a, and 4b in HEK293 cells after transfection of the cDNAs. Tubulin (orange), and RNF220 (green), DAPI (blue). Scale bar, 10μm.

In this presented study, we suggest that RNF220 isoform 4 plays a major role in mediating *RNF220* gene function in multiple tissues. Notably, isoform 4 is critical for muscle differentiation in myoblast cells and disappears upon terminal differentiation. Isoforms 1 and 4 may exhibit distinct subcellular localizations and subnuclear structures, with isoform 4 showing broader tissue expression than isoform 1. Given that the prior literature on RNF220 functions has been limited to analysis of the 566 aa isoform 1, our findings underscore the need to investigate the short isoform, isoform 4, to fully understand the biological functions of RNF220.

## Materials and Methods

### Mouse Tissues

The protocols of animal experiments were reviewed and approved by the Institutional Animal Care and Use Committee of Sookmyung Women’s University. Tissues were dissected from a 9-week-old male mouse and used for protein extraction followed by Western blotting.

### Cells

HEK293 cells were purchased from Korean Cell Line Bank and hESCs (H9) were purchased from Wicell Research Institute. C2C12s were obtained from American Type Culture Collection.

### Plasmids and siRNAs

Each isoforms’ EGFP-tagged RNF220 expression vector and non-tagged expression vector were cloned using hRNF220-WT (iso 1, kindly gifted by Cheol-Hee Kim and Seunghee Lee) (Kim, et al. 2018) and Q5 Site-Directed Mutagenesis Kit (NEB) according to the manufacturer’s protocol.

**Further information** is available in the **Supplementary Information**.

## Results

Four human RNF220 isoform polypeptides were annotated based on the RefSeq database (https://www.ncbi.nlm.nih.gov/refseq/). Detailed information on the transcriptional variants and their corresponding four peptide isoforms is summarized in **S Table 1**. Fifteen exons of the isoform 1 RNF220 transcript (NM_018150.4) were annotated with 2 alternative transcription start sites leading to isoforms initiating in exons 2a or 5a. The red indicated in **Fig. 1a** is not in the isoform 1 peptide and the yellow is only included in isoform 4. Interestingly, zebrafish and fruit fly express only a single, short isoform which corresponds to the human isoform 4 (**Fig. 1a, S Table 2**) suggesting that these long-overlooked short isoform of RNF220 may mediate the most conserved functions of this gene in evolution. We examined the expression pattern of RNF220 in mouse tissues using two antibodies, Ab1 and Ab2, that detect the N- and C-terminal regions respectively (**Fig. 1b, S Fig. 1**). Using Ab2 and less specific Ab1, we found that putative isoform 1 expresses only in the brain, whereas putative isoform 4 express in all the examined tissues. A human embryonic kidney cell line, HEK293, also expressed both isoform 1-like and isoform 4-like bands. These are depleted by siRNAs selectively targeting exons of each isoform (**Fig. 1c**). Based on molecular weight and siRNA-mediated suppression, we conclude that the isoforms primarily expressed in these cells and tissues are isoform 1 (63 kDa) and isoform 4 (35 kDa).

Using public ChIP-seq data (Carninci, et al. 2006; Forrest, et al. 2014), we sought to identify which RNF220 TSSs (transcription starts sites) for different isoforms might be utilized in various human tissues based on H3K4me3, which is a hallmark of active TSS chromatin (Guenther, et al. 2007). The peaks of H3K4me3 enrichment were observed that mapped to multiple RNF220 TSSs in all tissues examined **(S Fig. 2)**. The isoform 4 TSS showed a most highly enriched peak of H3K4me3 in all tissues examined except the hippocampus where the TSS for iso1/3 was stronger. These findings are consistent with the mouse immunoblotting results **(Fig. 1b)** and suggest that transcription of isoform 4 is more active than any other isoforms in most tissues, both in humans and mice.

**Figure 2.**
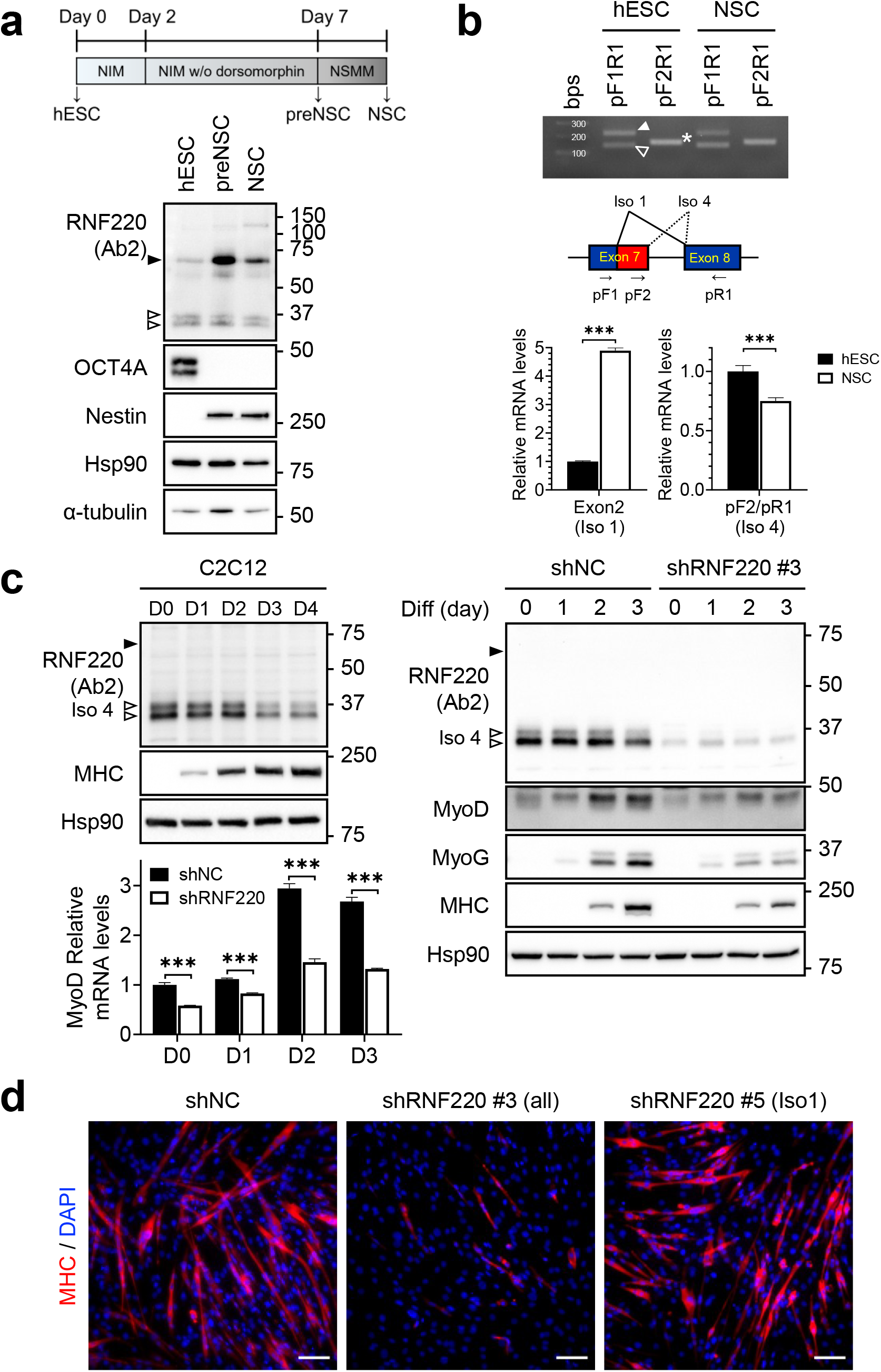
Expression of RNF220 isoforms during NSC differentiation and muscle differentiation. (a) Expression of the isoforms during NSC differentiation from hESC were tested with the all isoforms targeting Ab2. Only isoform 1 and 4 were detected. NIM : Neural Induction Medium, NSMM : Neural Stem Cell Maintenance Medium. (b) RT-PCR of exon7/8 alternative spliced mRNA for isoforms (pF1R1; 146 bp-iso1 and 224 bp-iso 4, pF2R1; 165 bp-iso 4). Isoform mRNA expression assessed by qRT-PCR using primers specific to exon 2 (isoform 1) and alternatively spliced exon 7 (pF2/pR1; isoform 4). Mean ± SD. ***, *P<0*.*001*. (c) Expression of RNF220 during C2C12 myoblast differentiation (left). Differentiation of C2C12 myoblasts, after transduced with lentivirus for control shRNA (shNC) or shRNF220 targeting all isoforms (#3), followed with the western blotting and qRT-PCR. (d) C2C12 transduced with lentivirus for control (shNC), targeting all isoforms (#3) or targeting isoform 1 only (#5) was differentiated. After 3days, cells were fixed and stained with anti-MHC and DAPI. Scale bar, 50 μm.

We noticed that the isoform 4 was detected as two bands. We found that the mRNA for isoform 4 may have a 2ndary start codon at M38, supported by another potential Kozak sequence which is conserved in evolution. This suggests the smaller band (4b) might be translated from this 2ndary start codon (**Fig. 1d**). Interestingly, using the 2ndary start codon may partially disrupt a putative nuclear localization signal. Overexpressed isoforms 1 and 4a are stained in nuclear and localized to nuclear speckle-like structures, forming a distinctive ring pattern, consistent with previous reports (Sferra, et al. 2021). In contrast, isoform 4b displayed a completely different nuclear staining pattern along with partial cytoplasmic distribution (**Fig. 1e**). Similar results were observed in live imaging of cells transfected with these EGFP-fused RNF220 isoforms (**S Fig. 3**). These findings suggest that isoform 4 exhibits distinguished biological roles from that of isoform 1 based on the subnuclear structure and subcellular localization.

Given the reported roles of RNF220 isoform 1, in murine neural stem cells (NSCs) (Ma et al. 2019; Zhang, et al. 2020) in mice, we analyzed the expression of RNF220 isoforms in NSCs derived from human embryonic stem cells (hESCs) (**Fig. 2a**). While both isoform 1 and isoform 4 expressed in hESCs, differentiation into NSCs resulted in increased isoform 1 and decreased isoform 4. A unique splicing pattern at exon 7 generated isoform 1, whereas other isoforms arose from alternative splicing (illustrated in **Figs. 1a** and **2b**). The changes of isoforms during hESCs differentiation into NSCs may be driven by isoform-specific transcription.

Notably, the muscle expressed only isoform 4 but less than other tissues (**Fig. 1b**). To investigate this further, we analyzed RNF220 expression during the differentiation of C2C12 murine myoblasts. Unexpectedly, cultured C2C12 cells strongly expressed isoform 4, but its expression decreased during differentiation into myotubes as expression of the sarcomere protein myosin heavy chain (MHC) increased (**Fig. 2c**). shRNA knockdown of all forms of RNF220 in C2C12 cells reduced expression of MyoD, the basic helix-loop-helix transcription factor responsible for muscle cell lineage determination, both before and during differentiation. RNF220 was required for the expression of both myogenic differentiation factor myogenin and MHC during differentiation. Reduced cell fusion to form myotubes and impaired differentiation when all RNF220 isoforms were targeted. In contrast, an shRNA specifically targets isoforms 1 failed to reveal the same effect (**Fig. 2d and S Fig. 4**). Overall, we conclude that it is isoform 4, rather than the full-length isoform 1, that is required for MyoD expression and subsequent differentiation into myotubes in C2C12 myoblasts. These findings further suggest that isoforms with the presence and absence of the N terminal half may not only exhibit tissue-specific expression but also perform distinct biological functions.

## Discussion

Recent studies have expanded the roles of RNF220 beyond neuronal tissues to other tissues and processes, such as cancer and immunity. In all of these studies, isoform 1 has been regarded as the primary functional molecule expressed from the *RNF220* locus, and the function was investigated. Similarly, *RNF220* knockout mice lacking exon 2, which eliminates isoform 1 but may not impact isoform 4, have been used in most studies to unveil the many novel biological functions of the RNF220 locus (except one study targeted exon 7; (Kim, et al. 2018)). Our findings reveal distinct expression patterns and functions for a short and N-terminally truncated RNF220, isoform 4. Isoform 4 is strongly expressed across most tissues, whereas isoform 1 is confined primarily to neuronal tissues in humans and mice. Isoform 4s appear crucial for myoblast differentiation. The dual translation initiation also modulates the subcellular localization of isoform 4a/b differently from isoform 1, underscoring their unique functional attributes. Familial leukodystrophy is caused by R363Q and R365Q mutations (Sferra, et al. 2021) affecting all isoforms. Neurological symptoms likely stem from abnormal isoform 1 due to its tissue-specific expression, while isoform 4, with broader expression, may drive systemic pathologies like cardiomyopathy. In skin fibroblasts, where isoform 1 mRNA was undetectable (Sferra, et al. 2021), mutant isoform 4a or 4b likely caused nuclear membrane abnormalities. The unique cytosolic and subnuclear structure of isoform 4b may further contribute to these phenotypes.

Isoform 4 is strongly expressed at the myoblast stage but disappears as myocyte differentiation progresses. The loss of RNF220 during differentiation suggests that non-dividing, terminally differentiated muscle cells do not express RNF220. For cells to maintain their differentiation potential as myoblasts do, functional RNF220 appears to be necessary.

The predominant expression of isoform 4 suggests that RNF220 may influence the differentiation of various tissues beyond the nervous system. The absence of long isoforms in zebrafish and fruit flies supports the idea that isoform 4 represents the original form of RNF220. Notably, this was also essential for nervous system patterning in zebrafish (Ma, Song, et al. 2020) and Drosophila (Sferra, et al. 2021), resembling the function of isoform 1 in humans.

Our results indicate that previous studies on RNF220, which primarily focused on isoform 1 expressed mostly in neuronal tissues, may have overlooked broader functional roles of isoform 4 across various tissues. Thus, future investigations should carefully consider that isoform 4 may play an important role in multiple tissues through its own function.

## Author Contributions

SC performed conceptualization and experiments, analyzed the data, wrote the manuscript and acquired funding. DJW performed experiment and analyzed data. JK and YHG performed experiment and provided samples. HJC and GUB provided expertise, resources and feedback. SJK provided expertise, wrote the manuscript and secured funding. WYK supervised the research, conceptualized, wrote the manuscript and secured funding.

## Acknowledgement

This study was supported by grants from the Korea Health Industry Development Institute (KHIDI), the National Research Foundation of Korea (NRF) funded by the Korean government (MSIT and MOE) [HI21C2509, 2022R1A5A2021216, 2020R1A2C1006091, RS-2024-00509503 and RS-2024-00463034]. DJW and SJK were supported by NIH R01s AG069865 and CA254047.

## Supplementary Materials and Methods

### Western Blotting

After tissue dissection from mouse, it was immediately frozen and ground with liquid nitrogen. Frozen tissues were lysed with ice-cold RIPA buffer containing protease and phosphatase inhibitors. Lysates were sonicated and centrifuged. Protein concentrations were determined using Pierce BCA Protein Assay kit (Thermo Fisher). After boiling in the sample buffer, protein lysates were separated on SDS-PAGE and transferred to PVDF membranes (Millipore-Sigma). Membranes were blocked with 5% skim milk in TBST and incubated overnight with primary antibodies in 5% BSA. Next, the membranes were incubated with HRP-conjugated secondary antibodies, the signals were detected using ECL substrate (Thermo Fisher) on the iBright imaging system (Thermo Fisher). Antibodies used for immunoblot analysis are as follows: Anti-RNF220 (Sigma-Aldrich, Ab1, HPA027578 and Ab2, HPA027577), Anti-α-tubulin (Santa Cruz Biotech, SC-23948), Anti-HSP 90α/β (F-8) (Santa Cruz Biotech, SC-13119), Anti-OCT4A (Cell Signaling Technology, 2840S), Anti-MHC (DSHB, MF20 (hybridoma)), Anti-MyoD (Santa Cruz Biotech, SC-377460), Anti-MyoG (DSHB, IF05 (hybridoma)).

### Cell culture

HEK293 cells were cultured in high glucose DMEM containing 10% fetal bovine serum, 100 units/ml penicillin-100 μl/ml streptomycin. hESCs were maintained in mTeSR1 or MACS-iPSC brew medium on plates coated with Matrigel diluted at 1:80 in DMEM/F12 for feeder-free conditions. The medium was replaced every day up to passaging, and the cells were enzymatically dissociated using a dispase solution. C2C12 myoblasts were cultured in the growth medium, high glucose DMEM containing 15% fetal bovine serum, 100 units/ml penicillin-100 μl/ml streptomycin. To induce myogenic differentiation, 80%-90% confluent C2C12 cells were exchanged into differentiation medium, high glucose DMEM containing 2% horse serum (Thermo Fisher) 100 units/ml penicillin -100 μl/ml streptomycin. All cells were maintained at 37°C under 5% CO2 in an incubator.

### TSS H3K4me3 enrichment analysis

The H3K4me3 ChIP-seq data for human tissues were obtained from the ENCODE portal (ENCODE Project Consortium. 2012; Luo, et al. 2020) (https://www.encodeproject.org/) with following identifiers: ENCFF504XAW, ENCFF231KHQ, ENCFF050YYY, ENCFF053IVC, ENCFF288QMN, ENCFF229BGF, ENCFF487CTD, ENCFF929KNV, ENCFF918PWP, ENCFF283ZMI and ENCFF237QAL. Fold change peaks for histone marks (H3K4me3) at RNF220 gene locus were used to each isoforms TSS enrichment analysis. TSS were annoted according to RefSeq mRNA in NCBI (https://www.ncbi.nlm.nih.gov/gene/55182).

### Confocal microscopy live imaging

For live imaging, HEK293 cells were transfected with EGFP-fused RNF220 isoform constructs for 24hrs. After transfection, cells were cultured in Leibovitz’s L-15 medium supplemented with 10% FBS and nuclei were counterstained with Hoechst 33342 (Thermo Scientific). The cells were placed on the plate that was heated to 37 °C and were observed using a LSM-700 (Zeiss) confocal microscope with 63x lens.

### Embryonic stem cells (ESC) and Neural stem cells (NSC)

To differentiation into NSC, human ESCs (hESCs, H9; Wicell Research Institute) cultured in StemMACSTM iPS-brew media (Miltenyi-Biotec) with Matrigel-coated plate transferred at approximately 20% confluence with 10 μM Y-27632. StemMACS media was then switched to Neural Induction Medium [NIM: 50% Advanced DMEM/F12, 50% Neurobasal Medium, 1% N2 supplement (100X), 2% B27 supplement minus vitamin A (50X), Glutamax, 10 ng/mL human LIF (R&D Systems), 4 μM CHIR99021 (Peprotech), 3 μM SB431542 (abcam), and 0.1 μM Compound E (Tocris)]. 2 μM of Dorsomorphin (Tocris) was added for two days and excluded for following five days. 7-days after differentiation, the cells were collected as preNSC to observe the change during differentiation to NSC or transferred into another matrigel-coated 35mm plate at a density of 400,000 cells, using the Accutase solution with Neural Stem Cell Maintenance Medium [NSMM: 50% Advanced DMEM/F12, 50% Neurobasal Medium, 1% N2 supplement (100X), 2% B27 supplement minus vitamin A (50X), Glutamax, and 10 ng/ml human LIF, 3 μM CHIR99021, 2 μM SB431542 containing 10 μM Y27632]. The NSCs were passaged every week using the Accutase solution.

### RNF220 knockdown using lentiviral shRNA

For shRNA lentivirus generation, pLKO.1-shNC (SHC002) or pLKO.1-shRNF220 (TRCN0000160783) (Sigma) was used. HEK293 cells were transfected with lentiviral packaging plasmids (psPAX2) and envelope expressing plasmids (pMD2.G) (Addgene) using jetPRIME reagent. After transfection, the supernatant containing lentivirus was collected and filtered (0.45μm). The aliquots were stored at -80°C. C2C12 cells were infected using 8 μg/ml polybrene and selected with 2 μg/ml puromycin for 3 days.

### Quantitative RT-PCR analysis

Total RNA was extracted from cells using Trizol (Invitrogen) according to the manufacturer’s instructions, then cDNA was synthesized by iScript reverse transcriptase (Bio-Rad). qRT-PCR analysis performed with Power SYBR green (Applied Biosystems). The relative mRNA levels of each gene or transcript variants were normalized to the level of GAPDH or β-actin. Primers are listed below. RNF220 5’-GCA GCC TTC AAG ATG GAG AAC-3’ (Exon2-F), 5’-GAG ACA GGT ACA CCA AAG GGT-3’ (Exon2-R), 5’-GAG GAA GCA AGA TGA AGG GC-3’ (pF1), 5’-AGC AGG AGA TGA GTA GGC ATG-3’ (pF2) and 5’-CAC CTT CCA GGA GAG TGG TG-3’ (pR1); β-actin 5’-ATT GGC AAT GAG CGG TTC-3’ (β-actin-F) and 5’-GGA TGC CAC AGG ACT CCA T-3’ (β-actin-R); mouse GAPDH 5’-CTT TGT CAA GCT CAT TTC CTG G-3’ (mGAPDH-F) and 5’-TCT TGC TCA GTG TCC TTG C-3’ (mGAPDH-R); mouse MyoD 5’-GCG GTT CAG GAC CAC TTA TT-3’ (mMyoD-F) and 5’-CAG CAT GCC TGG GAG ATA AA-3’ (mMyoD-R).

### Immunofluorescence staining

For immunostaining for cells, cells were fixed using 4% paraformaldehyde and permeabilized using 0.1% triton X-100. After blocking with 5% goat serum (Thermo Fisher) or 5% BSA, they were incubated with primary antibodies and fluorochrome-linked secondary antibodies. Nuclei were counterstained with DAPI. Images were acquired by LSM-700 (Zeiss), and analyzed by using ImageJ or ZEISS Zen Imaging Software.

### Statistical analysis

All data were presented as mean ± standard deviation (SD), as indicated in the figure legends. GraphPad Prism 8 software was utilized for all statistical analyses. For comparisons between the two groups, statistical significance was evaluated using an unpaired t-test. Statistical significance was indicated as ***, P <0.001.

**S Figure 1.**
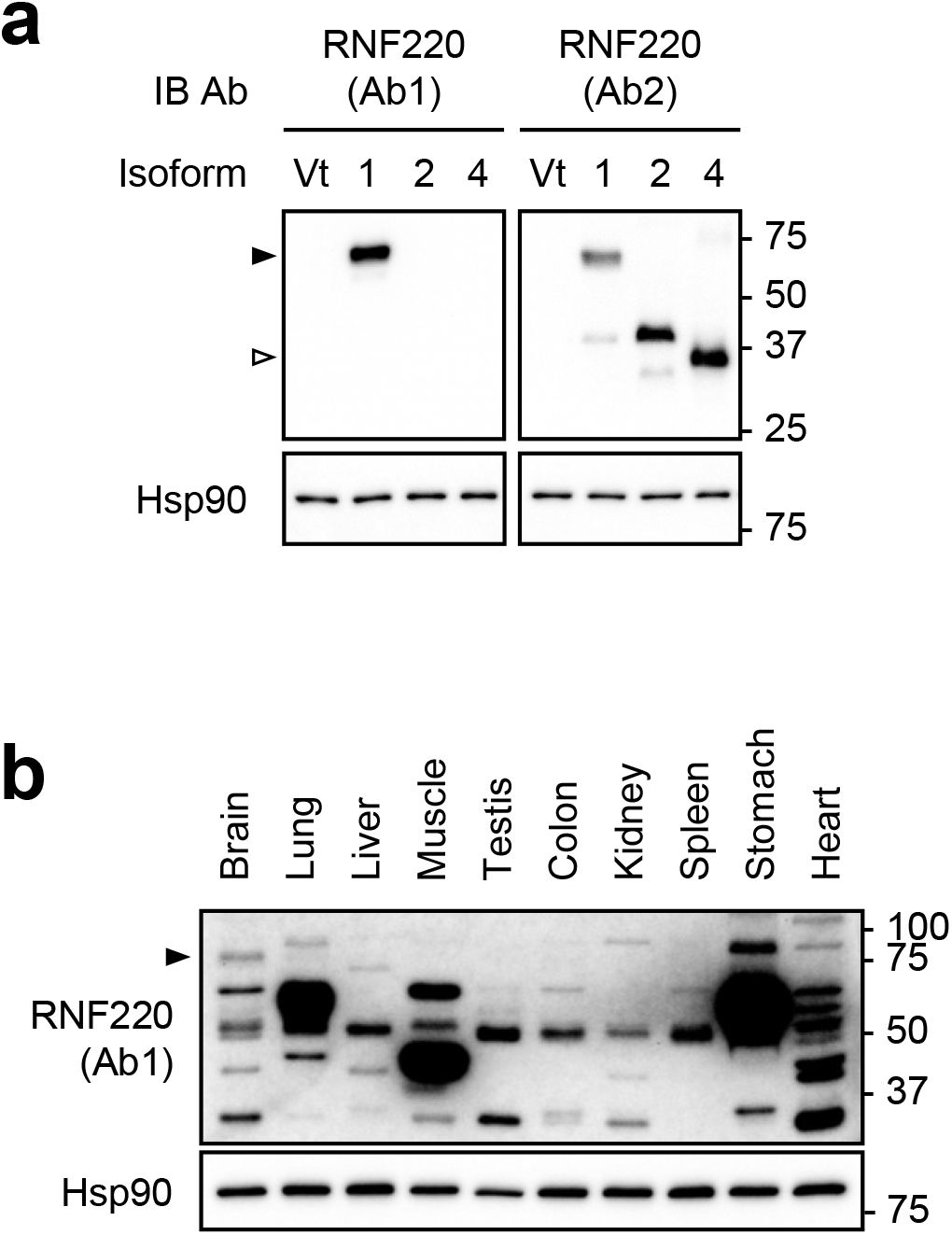
The expression of isoform 1 limited to the brain tissue in mouse. (a) Re-validation of two anti-RNF220 affinity-purified rabbit polyclonal antibodies, Ab1 and Ab2 (Atlas Antibodies HPA027578 and HPA027577), which were generated against recombinant PrEST antigens encoding the N-terminal (aa 44–145) and C-terminal (aa 375–454) regions, respectively. Overexpression of cDNA for each isoform in HEK293 cells conforms to the specificity of the Abs. (b) Western blot analysis of mouse tissues using Ab1. Despite Ab1 generating multiple nonspecific bands, isoform 1 is detected exclusively in the brain, as shown in Figure 1b in the main manuscript.

**S Figure 2.**
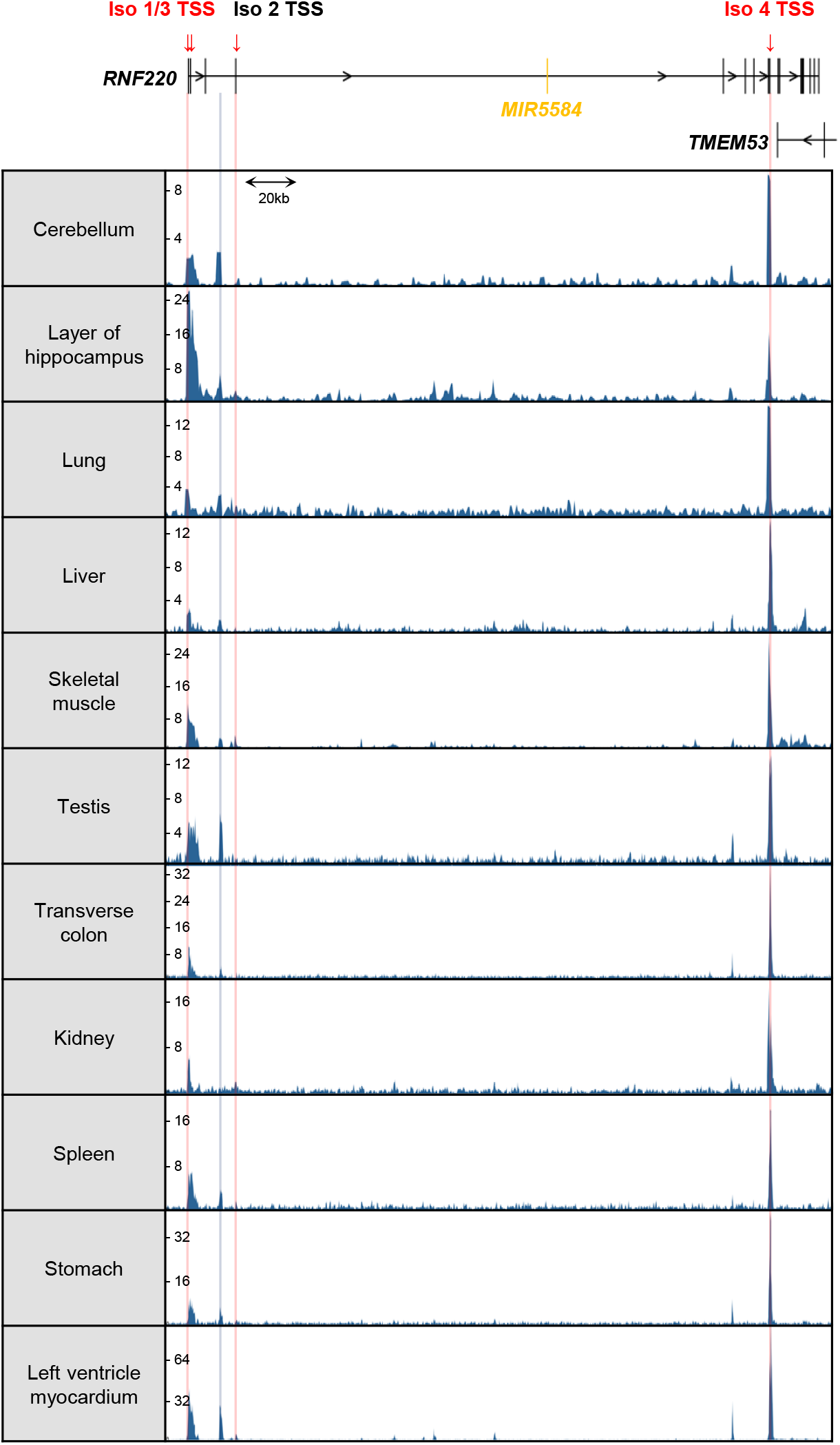
Isoform 4 TSS region was more enriched with H3K4me3 than isoform 1 TSS in most human tissue. Genome browser snapshot of H3K4me3 enrichment at RNF220 gene locus in various human tissues. The published H3K4me3 ChIP-seq data are used (ENCODE Project Consortium. 2012; Luo, et al. 2020). Red lines mark each isoform transcription start site.

**S Figure 3.**
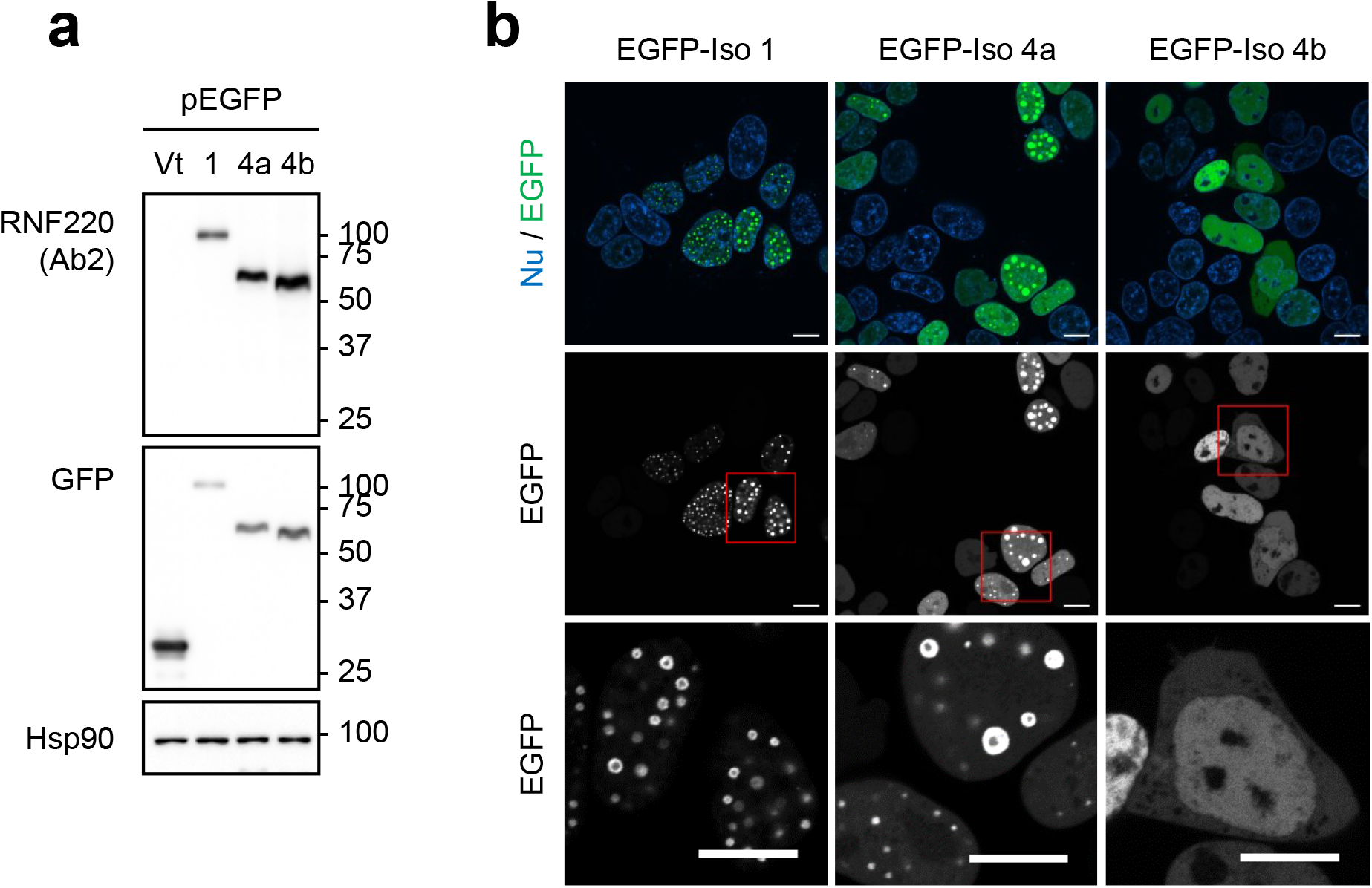
RNF220 isoform 1 and 4a make distinct structure in the nucleus. (a) Western blot analysis of EGFP-RNF220 isoforms expressed in HEK293 cells. (b) In vivo live imaging of EGFP-tagged RNF220 isoforms (iso 1, iso 4a, and iso 4b). The constructs were introduced into HEK293 cells, and 24 hours post-transfection, nuclei were stained with Hoechst 33342 (Nucleus, NU), followed by live imaging using confocal microscopy. The third row shows magnified images of the insets in the second row. The unique cytosolic localization and loss of the ring-like structure of isoform 4b, distinguishing it from isoform 1 and isoform 4a, are evident. Scale bar, 10μm.

**S Figure 4.**
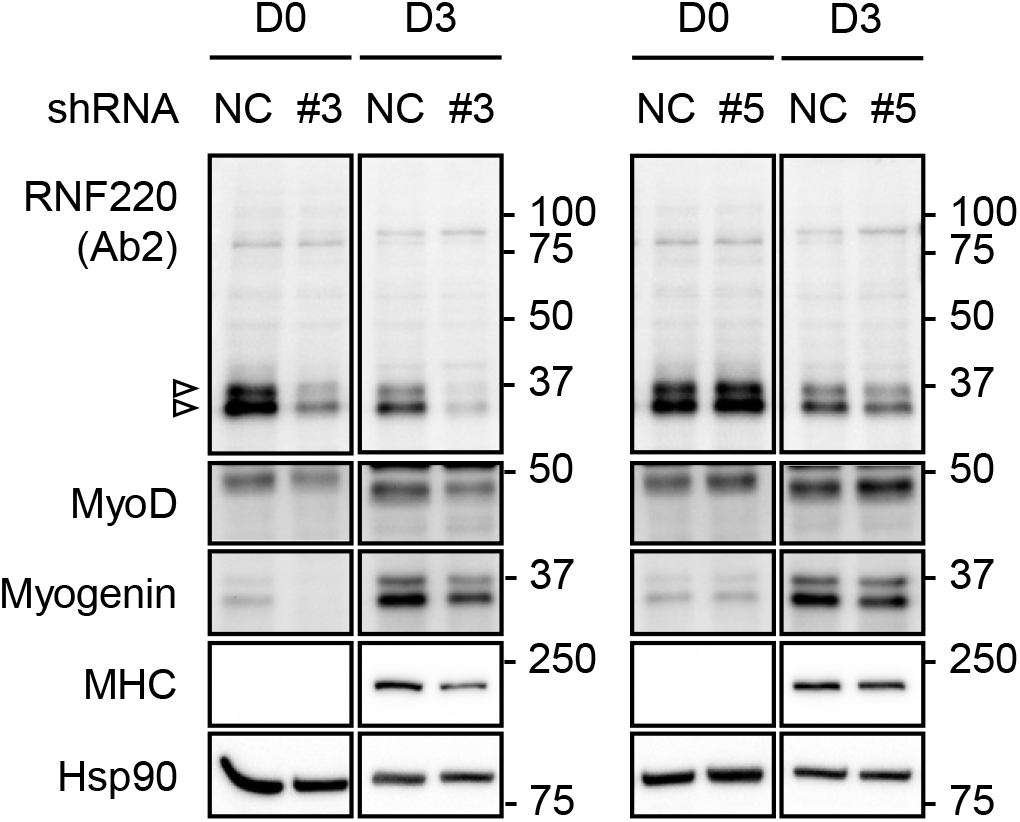
Not the RNF220 isoform 1 but the isoform 4 is required for myogenic differentiation. Western blot for MyoD, Myogenin and MHC in C2C12 myoblast differentiation at day 0 and day 3 after the C2C12 is transduced with lentivirus for control shRNA or shRNF220 targeting all isoforms (#3) or isoform 1 (#5).

**S Table 1.** Summary of the RNF220 isoforms curated with RefSeq mRNA in NCBI (https://www.ncbi.nlm.nih.gov/gene/55182)

| Isoforms | Isoform 1 |  |  | Isoform 2 | Isoform 3 |  | Isoform 4 |
| --- | --- | --- | --- | --- | --- | --- | --- |
| BLAST mRNA | *NM_018150.4* | NM_001319956.1 | NM_001376488.1 | *NM_001319957.2* | *NM_001376486.1* | NM_001376487.1 | *NM_001376489.1* |
| RNA title | *transcript variant 1* | transcript variant 2 | transcript variant 6 | *transcript variant 3* | *transcript variant 4* | transcript variant 5 | *transcript variant 7* |
| BLAST protein | *NP_060620.2* | NP_001306885.1 | NP_001363417.1 | *NP_001306886.1* | *NP_001363415.1* | NP_001363416.1 | *NP_001363418.1* |
| TXN start | **44,405,255** | 44,405,160 | 44,406,792 | 44,424,001 | **44,405,255** | **44,405,255** | 44,631,968 |
| TXN stop | **44,651,723** | 44,651,724 | **44,651,723** | **44,651,723** | **44,651,723** |  | **44,651,723** |
| TXN Span | **246,469 nt** | 246,565 nt | 244,932 nt | 227,723 nt | **246,469 nt** |  | 19,756 nt |
| mRNA length | 3,042 nt | 3,134 nt | 3,056 nt | 2,241 nt | 3,120 nt | 3,116 nt | 2,062 nt |
| CDS length | 1,701 nt |  |  | 1,062 nt | 1,779 nt |  | 927 nt |
| Protein length | 566 aa |  |  | 353 aa | 592 aa |  | 308 aa |
| Protein size | 62.8 kDa |  |  | 39.6 kDa | 65.7 kDa |  | 34.7 kDa |

*Italic* : Used as a reference for each isoform in this paper.
**Bold** : Same as other isoforms.

**S Table 2.** Amino acid sequence similarity (in percentage) of RNF220 isoforms, among human, mouse, chicken, zebrafish and fruit fly.

|  | Mus musculus<br>house mouse |  |  |  | Gallus gallus<br>chicken |  | Danio rerio<br>zebrafish | Drosophila<br>melanogaster<br>fruit fly |
| --- | --- | --- | --- | --- | --- | --- | --- | --- |
| Symbol | RNF220 |  |  |  | RNF220 |  | rnf220a | RNF220 |
| Genomic context_Location | ch4 |  |  |  | ch8 |  | ch2 | ch3 |
| Isoform | isoform 1 | isoform 2 | isoform 3 | isoform 4 | isoform 1 | isoform 2 | - | - |
| aa # | 566 | 353 | 308 | 245 | 305 | 589 | 307 | 398 |
| RefSeq accession number | NP_080015.3 | NP_001297658.1 | NP_001297659.1 | NP_001297660.1 | NP_001012617.1 | NP_001103457.1 | NP_001070038.1 | NP_651088.1 |
| vs human isoform 1 (566aa) | 98% | 54% | 45% | 42% | 44% | 90% | 37% | 25% |
| vs human isoform 2 (592aa) | 54% | 97% | 82% | 68% | 80% | 56% | 69% | 33% |
| vs human isoform 3 (353aa) | 94% | 58% | 49% | 40% | 48% | 94% | 41% | 24% |
| vs human isoform 4 (308aa) | 45% | 81% | 97% | 78% | 94% | 48% | 78% | 32% |

Similarity is calculated by using Needleman-Wunsch alignment.

## REFERENCES

Aigner P, Just V, Stoiber D (2019) STAT3 isoforms: Alternative fates in cancer? Cytokine 118: 27–34. doi: 10.1016/j.cyto.2018.07.014

Carninci P, Sandelin A, Lenhard B, Katayama S, Shimokawa K, Ponjavic J, Semple CA, Taylor MS, Engström PG, Frith MC (2006) Genome-wide analysis of mammalian promoter architecture and evolution. Nat Genet 38: 626–635

Davuluri RV, Suzuki Y, Sugano S, Plass C, Huang TH (2008) The functional consequences of alternative promoter use in mammalian genomes. Trends Genet 24: 167–177. doi: 10.1016/j.tig.2008.01.008

de Klerk E, ‘t Hoen PAC (2015) Alternative mRNA transcription, processing, and translation: insights from RNA sequencing. Trends Genet 31: 128–139. doi: 10.1016/j.tig.2015.01.001

Deng T, Zhong P, Lou R, Yang X (2023) RNF220 promotes gastric cancer growth and stemness via modulating the USP22/wnt/beta-catenin pathway. Tissue Cell 83: 102123. doi: 10.1016/j.tice.2023.102123

Forrest ARR, Kawaji H, Rehli M, Baillie JK, de Hoon MJL, Haberle V, Lassmann T, Kulakovskiy IV, Lizio M, Itoh M, Andersson R, Mungall CJ, Meehan TF, Schmeier S, Bertin N, Jørgensen M, Dimont E, Arner E, Schmidl C, Schaefer U, Medvedeva YA, Plessy C, Vitezic M, Severin J, Semple CA, Ishizu Y, Young RS, Francescatto M, Alam I, Albanese D, Altschuler GM, Arakawa T, Archer JAC, Arner P, Babina M, Rennie S, Balwierz PJ, Beckhouse AG, Pradhan-Bhatt S, Blake JA, Blumenthal A, Bodega B, Bonetti A, Briggs J, Brombacher F, Burroughs AM, Califano A, Cannistraci CV, Carbajo D, Chen Y, Chierici M, Ciani Y, Clevers HC, Dalla E, Davis CA, Detmar M, Diehl AD, Dohi T, Drabløs F, Edge ASB, Edinger M, Ekwall K, Endoh M, Enomoto H, Fagiolini M, Fairbairn L, Fang H, Farach-Carson MC, Faulkner GJ, Favorov AV, Fisher ME, Frith MC, Fujita R, Fukuda S, Furlanello C, Furino M, Furusawa J, Geijtenbeek TB, Gibson AP, Gingeras T, Goldowitz D, Gough J, Guhl S, Guler R, Gustincich S, Ha TJ, Hamaguchi M, Hara M, Harbers M, Harshbarger J, Hasegawa A, Hasegawa Y, Hashimoto T, Herlyn M, Hitchens KJ, Ho Sui SJ, Hofmann OM, Hoof I, Hori F, Huminiecki L (2014) A promoter-level mammalian expression atlas. Nature 507: 462–470. doi: 10.1038/nature13182

Gleason AC, Ghadge G, Sonobe Y, Roos RP (2022) Kozak similarity score algorithm identifies alternative translation initiation codons implicated in cancers. International Journal of Molecular Sciences 23: 10564

Guenther MG, Levine SS, Boyer LA, Jaenisch R, Young RA (2007) A chromatin landmark and transcription initiation at most promoters in human cells. Cell 130: 77–88

Guo X, Ma P, Li Y, Yang Y, Wang C, Xu T, Wang H, Li C, Mao B, Qi X (2021) RNF220 mediates K63-linked polyubiquitination of STAT1 and promotes host defense. Cell Death Differ 28: 640–656. doi: 10.1038/s41418-020-00609-7

Kim J, Choi T, Park S, Kim MH, Kim C, Lee S (2018) Rnf220 cooperates with Zc4h2 to specify spinal progenitor domains. Development 145: dev165340. doi: 10.1242/dev.165340. doi: 10.1242/dev.165340

Kosugi S, Hasebe M, Tomita M, Yanagawa H (2009) Systematic identification of cell cycle-dependent yeast nucleocytoplasmic shuttling proteins by prediction of composite motifs. Proceedings of the National Academy of Sciences 106: 10171–10176

Kubickova A, De Sanctis JB, Hajduch M (2023) Isoform-Directed Control of c-Myc Functions: Understanding the Balance from Proliferation to Growth Arrest. Int J Mol Sci 24: 17524. doi: 10.3390/ijms242417524. doi: 10.3390/ijms242417524

Ma P, An T, Zhu L, Zhang L, Wang H, Ren B, Sun B, Zhou X, Li Y, Mao B (2020) RNF220 is required for cerebellum development and regulates medulloblastoma progression through epigenetic modulation of Shh signaling. Development 147. doi: 10.1242/dev.188078

Ma P, Song N, Cheng X, Zhu L, Zhang Q, Zhang LL, Yang X, Wang H, Kong Q, Shi D, Ding Y, Mao B (2020) ZC4H2 stabilizes RNF220 to pattern ventral spinal cord through modulating Shh/Gli signaling. J Mol Cell Biol 12: 337–344. doi: 10.1093/jmcb/mjz087

Ma P, Song N, Li Y, Zhang Q, Zhang L, Zhang L, Kong Q, Ma L, Yang X, Ren B, Li C, Zhao X, Li Y, Xu Y, Gao X, Ding Y, Mao B (2019) Fine-Tuning of Shh/Gli Signaling Gradient by Non-proteolytic Ubiquitination during Neural Patterning. Cell Reports 28: 541. doi: 10.1016/j.celrep.2019.06.017

Ma P, Wan LP, Li Y, He C, Song N, Zhao S, Wang H, Ding Y, Mao B, Sheng N (2022) RNF220 is an E3 ubiquitin ligase for AMPA receptors to regulate synaptic transmission. Sci Adv 8. doi: 10.1126/sciadv.abq4736

Pan Y, An N, Deng X, Zhang Q, Du X (2021) RNF220 promotes the proliferation of leukaemic cells and reduces the degradation of the Cyclin D1 protein through USP22. Blood Cells Mol Dis 86: 102490. doi: 10.1016/j.bcmd.2020.102490

Sferra A, Fortugno P, Motta M, Aiello C, Petrini S, Ciolfi A, Cipressa F, Moroni I, Leuzzi V, Pieroni L, Marini F, Boespflug Tanguy O, Eymard-Pierre E, Danti FR, Compagnucci C, Zambruno G, Brusco A, Santorelli FM, Chiapparini L, Francalanci P, Loizzo AL, Tartaglia M, Cestra G, Bertini E (2021) Biallelic mutations in RNF220 cause laminopathies featuring leukodystrophy, ataxia and deafness. Brain 144: 3020–3035. doi: 10.1093/brain/awab185

Song N, Ma P, Zhang Q, Zhang L, Wang H, Zhang L, Zhu L, He C, Mao B, Ding Y (2020) Rnf220/Zc4h2-mediated monoubiquitylation of Phox2 is required for noradrenergic neuron development. Development 147: dev185199. doi: 10.1242/dev.185199. doi: 10.1242/dev.185199

Wang Y, Song N, Zhang L, Ma P, Chen J, Huang Y, Hu L, Mao B, Ding Y (2022) Rnf220 is Implicated in the Dorsoventral Patterning of the Hindbrain Neural Tube in Mice. Front Cell Dev Biol 10: 831365. doi: 10.3389/fcell.2022.831365

Yan J, Tan M, Yu L, Jin X, Li Y (2021) Ring finger 220 promotes the stemness and progression of colon cancer cells via Ubiquitin specific peptidase 22-BMI1 axis. Bioengineered 12: 12060–12069. doi: 10.1080/21655979.2021.2003664

Zhang L, Ye M, Zhu L, Cha J, Li C, Yao Y, Mao B (2020) Loss of ZC4H2 and RNF220 Inhibits Neural Stem Cell Proliferation and Promotes Neuronal Differentiation. Cells 9: 1600. doi: 10.3390/cells9071600. doi: 10.3390/cells9071600

